# Evaluation of Female Sexual Function in Persons with Type 2 Diabetes Mellitus Seen in a Tertiary Hospital in South East Nigeria with Emphasis on Its Frequency and Predictors

**DOI:** 10.1101/526301

**Authors:** U Ezeani Ignatius, U Onyeonoro Ugochukwu, E UgwuTheophilus

**Affiliations:** Division of Endocrinology, Diabetes and Metabolism, Department of Internal Medicine, Federal Medical Center, Umuahia, Abia state, Nigeria; Department of Community Medicine, Federal Medical Center, Umuahia, Abia state, Nigeria; Department of Internal Medicine, Enugu State University Teaching Hospital, Enugu, Nigeria

**Keywords:** Female Sexual function, Diabetes mellitus, Frequency, Predictors, South east Nigeria, Dysfunction

## Abstract

**Background:** women with diabetes are at increased risk of sexual problems, however, this problem is under reported hence the need for this study.

**Methods:** This was a cross sectional case-controlled study. Seventy-five consenting females with type 2 DM were enrolled from the Diabetes Clinic of the Federal Medical Center, Umuahia, while Seventy-five persons which included hospital workers and female companions of subjects were recruited as control. Sexual dysfunction in both groups was diagnosed and characterized using the female sexual function index (FSFI). Data obtained from this study was presented as Mean±SD and analyzed using SPSS 17 software.

**Results:** The mean age of the T2DM group and control were 44.5 years and 38.9 years respectively. The mean total female sexual score (TFSS) was 22.10±6.66 in the T2DM subjects, while in the control subjects, it was 22.43±5.29. This was not statistically significant. The FSF scores in the desire, lubrication and orgasm domains were all lower in the diabetic women and this was statistically significant (P< 0.05). The domains of pain and arousal were also lower in the diabetic women although this was not statistically significant (P >0.05). The proportion of diabetic females who reported problems in the arousal, lubrication, orgasm and pain domains were higher (40.0, 36.4, 32.7, 29.1) than the controls (27.9, 16.2, 14.7, 19.1) {p<0.05}.

**Conclusion:** The prevalence of female sexual dysfunction was high from our study. Similarly, the Female Sexual Function Index (FSFI) score was low in women with diabetes when compared with controls. The domains of arousal, pain, orgasm and satisfaction were the most affected domains in subjects with DM Age, marital status, BMI, FBS and hypertension were predictive of sexual dysfunction in the diabetic women.

## 1. Introduction

Diabetes Mellitus occurs throughout the world. According to the International Diabetes Federation (IDF) eighth atlas, about 425 million people worldwide, or 8.8% of adults 20-79 years, are estimated to be living with diabetes mellitus in 2017.^1^ There is a relationship between diabetes and sexual dysfunction (SD): this has been noticed in both male and female.^2,3,4^Sexual dysfunctions in women with diabetes mellitus are often under reported when compared with men with diabetes. To the best of our knowledge, there are few studies in our environment that have focused on female sexual dysfunction, even though more cases are seen in the outpatient clinics than the number reported if any. Some probable reasons for this observation includes: 1. Women are still viewed as sexual objects in some societies and as a result of this, they are expected to accept sex and sexuality as a prelude for conception. Secondly, some societies view women who raise the issue of their sexual dysfunctions as promiscuous, this inadvertently will make them to conceal these challenges for fear of societal ridicule. In the early nineteenth century, before the discovery of insulin, sexuality was not a common topic of discourse neither was it an area that had benefited from extensive research. The initially conceived idea about sexual dysfunction in both sexes was, “If you do not ask about it, it does not exist.” The connection between diabetes and sexual function only began to be highlighted about a century ago unfortunately; more attention was given to male dysfunction. Furthermore, most of the publications placed emphasis on the effect of diabetes on male sexual function, not until the famous reproductive endocrinologist: Robert Kolodny reported the relationship between diabetes and female sexual dysfunction.^5^ There are several causes of female SD and these includes: vascular, neurological, endocrine and psychogenic causes, all these factors have been identified in the aetiology of female sexual dysfunction.^6^ Unlike male SD, female SD is majorly influenced by psychogenic factors such as depression whose occurrence is more than double in women when compared to their male counterparts.^6^

The probability of a woman with diabetes developing sexual dysfunction is higher when compared with those without DM. Sexual problems in women with diabetes could present in various ways. Some of these problems include dyspareunia, inadequate vaginal lubrication reduced arousal and desire. Even though there are studies on this subject from other parts of the world, literature on this subject from Nigeria is scarce, hence the need for this study.

### 1.1 Aims

The aim of this study is to examine the prevalence of sexual dysfunction in women with type 2 diabetes mellitus, compare the prevalence of sexual dysfunction in women with diabetes to that of a control group and describe the predictors of sexual dysfunction in women with diabetes.

## 2. Methodology

This was a cross sectional case-controlled study. Seventy-five consenting females with type 2 DM were enrolled from the Diabetes Clinic of the Federal Medical Center, Umuahia, Abia state. The inclusion criteria include subjects married for atleast 1year and have had a stable marital relationship. Patients who were on drugs like beta blockers and centrally acting drugs like alpha methyldopa known to cause female SD were excluded. Seventy-five persons which included hospital workers and female companions of subjects were recruited as control (these subjects were screened for diabetes). The questionnaire was administered by both male and female medical personnel in the diabetic unit who informed the subjects about the research and its objectives and they were assured that confidentiality will be maintained during and after the study. Information given was used only for the purpose of this study. All the staff working for the study were trained and examined before the enrollment. Information obtained from study and control subjects included age, marital status, educational status, employment history, drug history, type and duration of DM, height, weight, body mass index, waist circumference, hip circumference, and blood pressure. The weight obtained was recorded in kilograms (kg) to the nearest 0.1kg and the height recorded in meters (m) to the nearest 0.01m. The body mass index was calculated as the weight in kg divided by the square of the height in metres.^7^The waist circumference was measured using a non-stretch metric tape and taken at the mid-point between the rib cage and iliac crest while hip circumference was taken as the maximal circumference of the buttocks.^8^

Sexual dysfunction in both groups was diagnosed and characterized using the female sexual function index (FSFI)^9^ which is a specific, sensitive and standardized tool for diagnosing female SD. The index is a 19-item questionnaire providing scores on six domains of sexual function (desire, arousal, lubrication, orgasm, satisfaction, and pain) as well as a total score.^9,10,11^In women, the minimum and maximum scores are respectively 2 and 36. Women with a score under 26 were classified as having sexual dysfunction. This cut-off point was the same figure validated by other researchers. It is a well-accepted self-report instrument for assessing sexual function of women world-wide. The data obtained from this study was presented as Mean±SD and analyzed using SPSS 17 software.

## 3. Results

Between October 2016 and September 2017, 150 married women were studied (seventy-five diabetic women and seventy-five controls), but one hundred and twenty three returned there questionnaire. They were grouped into a diabetic group (*n=55*) and a non-diabetic group (*n=68*). Women with diabetes mellitus were those attending the Diabetes and Endocrinology clinics at the Federal Medical Center, Umuahia, Abia state and non-diabetic women were their female companions and health workers at the medical center. The mean age of the T2DM group and control were 44.5 years and 38.9 years respectively. This was statistically significant (p=0.04, Table 2). The proportion of persons who had some form of education was higher in the control subjects than in patients with T2DM and this was statistically significant (p=0.02).A greater majority of the control subjects were either self-employed or civil servants compared with the subjects with T2DM, although this was not statistically significant (p=0.24). A higher proportion of the control subjects were either overweight or obese when compared with subjects with T2DM, this was not statistically significant (p=0.33). The prevalence of SD in this study was 79.2% and the mean age was 47.3±7.9. The proportion of diabetic females who reported problems in the arousal, lubrication, orgasm and pain domains was 40.0, 36.4, 32.7 and 29.1 respectively. On the other hand the proportion in the control was 27.9, 16.2, 14.7 and 19.1 respectively. Age, marital status, BMI, FBS and hypertension are predictive of sexual dysfunction in the diabetic women (OR: 1.80, 1.15, 1.67, 1.00, 8.51).

## 4. Discussion

Sexual dysfunction (SD) is known to be common in male and females with DM, although it is grossly under reported in females with DM. The prevalence of female sexual dysfunction (FSD) in this study was 29.1%. This is much higher than the 6.6% reported by Unadike et al^12^ though it is almost same as the prevalence reported by Enzlinet al^13^ in the population they studied. Although the study by Unadike et al was performed in a region with a the same financial and educational background as ours, the low prevalence he reported may be as a result of changing perceptions by women (as a result of increasing modernization) on issues bordering on sexual challenges considering the fact that his study was carried out almost a decade ago. Women are becoming increasingly more informed and confident in expressing their opinions: this could be responsible for obvious increase in prevalence. Other studies reported even higher prevalence compared to findings in this study.^14,15^The complications of diabetes seem to have a much bigger influence on sexual problems as noted in our study.

The mean (SD) ages of subjects with T2DM were higher than that of the controls and this was statistically significant: increasing age was associated with the development of FSD. In studies from other countries, the age of the study population may have affected the FSD prevalence in such climes; a Nigerian study had much older subjects^16^ while a Belgium study enrolled the youngest participants.^13^ In our study, both the prevalence and age were moderate, similar to what was reported in a US study. Age has a significant impact on the sexual function of a woman as increasing age may be associated with declining sexual interest. With aging, women tend to experience hormonal changes such as estrogen/androgen reduction, which frequently cause significant bodily and emotional unpleasant effects on sexual function.^17^ This could explain the reason behind the varying prevalence rates of FSD noted in different studies.

The mean total female sexual function index (FSFI) score in T2DM subjects and their control were 22.1 and 22.4 respectively (p>0.05): this is in keeping with reports from other studies.^18,19,20^The FSF scores in the desire, lubrication and orgasm domains were all lower in the diabetic women and this was statistically significant (P< 0.05). The domains of pain and arousal were also lower in the diabetic women although this was not statistically significant(P >0.05). In the diabetic women, majority of subjects reported problems in the domains of arousal, lubrication, orgasm, satisfaction and pain when compared to the control group. This finding is in keeping with results from a study by Olarinoye et al^21^ who in a study involving fifty one type 2 DM women, noted arousal, pain, orgasm and satisfaction as the most affected domains.

In absolute percentage, the proportion of diabetic females who reported problems in the arousal, lubrication, orgasm and pain domains were higher than the controls. These differences were statistically significant in the two domains of orgasm and lubrication (p < 0.05). This value is higher than results of a Malaysian study.^22^ This difference could be attributed to the difference in culture, ideologies and religion: system of secularism in South East Nigeria with large inhabitants of Christians as compared with a predominantly Muslim population in the Malaysian study. This will influence expression of sexual opinions and thoughts and inexorably, cause the women to suppress topics relating to their sexuality for fear of its negative perception from the society. Thus, these sexual problems may go unreported.

Age, marital status, BMI, FBS and hypertension are predictive of sexual dysfunction in the diabetic women. Higher BMI class is predictive of sexual dysfunction in the diabetic women: this finding is similar to reports from a New York study.^23^In a study done in China, similar trend was reported although this was not seen in the non diabetic control group. Although study comparison between nations is problematic because varying definition and research methods were employed in these various studies. Another interesting finding from this study is the lower BMI and difference in HC and WC in patients with diabetes when compared to the control group. A possible explanation could be that in a patient with diabetes, a vital aspect of management is lifestyle intervention with one goal being weight reduction. Therefore, it may not be uncommon to see patients with T2DM having a lower BMI, difference in HC and WC. We feel that there is need for more studies to further investigate the mechanisms of obesity and sexual dysfunction in diabetic women.

The strength of our study lies in the use of the FSFI questionnaire, a validated instrument to assess female sexual function which has been extensively used in studies. Limitations that arose from this study include: This was a small study which should be considered exploratory, no multiple comparison adjustments were made in the analysis; therefore a larger and specifically designed study is needed to evaluate other clinical and metabolic abnormalities in patients with SD Secondly, we did not consider sex hormones, history of reproductive system diseases and other factors in this study.

## 5. Conclusion

The prevalence of female sexual dysfunction was high from our study. Similarly, the Female Sexual Function Index (FSFI) score was low in women with diabetes when compared with controls. The domains of arousal, pain, orgasm and satisfaction were the most affected domains in subjects with DM Age, marital status, BMI, FBS and hypertension were predictive of sexual dysfunction in the diabetic women. There may be need for more research to look at the influence of diabetes type on sexual function in order to explore various treatment strategies for this group of women.

## Consent statement

Written informed consent was obtained from the patient for publication of this research article. A copy of the written consent is available for review by the Editor-in-Chief of this journal.

## Declarations

Ethical approval: The Ethics and Research committee of the Federal Medical Center, Umuahia gave the ethical approval. The patients interviewed in this study did it voluntarily, and wrote an informed consent.

Source of funding: none

## Authors contribution

EI conceived of the study, carried out the sequence alignment and drafted the manuscript. OU and TU participated in the sequence alignment, design of the study and helped to draft the manuscript. All authors read and approved the final manuscript.

## Acknowledgements

We thank all the staff in the department of Internal Medicine Federal Medical Center, Umuahia. who contributed towards the article by making substantial contributions to conception and revision of manuscript for important intellectual content.

## Appendix I Female Sexual Function Index (FSFI) □

SubjectIdentifier________

Date_______

INSTRUCTIONS: These questions ask about your sexual feelings and responses <u>during the past 4weeks</u>. Please answer the following questions as honestly and clearly as possible. Your responses will be kept completely confidential. In answering these questions the following definitions apply:

<u>Sexual activity</u> can include caressing, foreplay, masturbation and vaginal intercourse.

<u>Sexual intercourse</u> is defined as penile penetration (entry) of the vagina.

<u>Sexual stimulation</u> includes situations like foreplay with a partner, self-stimulation (masturbation), or sexual fantasy.

### CHECK ONLY ONE BOX PER QUESTION

<u>Sexual desire</u> or <u>interest</u> is a feeling that includes wanting to have a sexual experience, feeling receptive to a partner’s sexual initiation, and thinking or fantasizing about having sex.

1. Over the past 4weeks, how **often** did you feel sexual desire or interest?
  □ Almost always or always
  □ Most times (more than half the time)
  □ Sometimes (about half the time)
  □ A few times(less than half the time)
  □ Almost never or never
2. Over the past 4weeks, how would you rate your **level** (degree) of sexual desire or interest?
  □ Very high
  □
  □
  □
  □
  High
  Moderate
  Low
  Very low
  or none
  Sexual arousal is a feeling that includes both physical and mental aspects of sexual excitement. It may include feelings of warmth or tingling in the genitals, lubrication (wetness),or muscle contractions.
3. Over the past 4weeks, how **often** did you feel sexually aroused (“turned on”) During sexual activity or intercourse?
  □ No sexual activity
  □ Almost always or always
  □ Most times (more than half the time)
  □ Sometimes (about half the time)
  □ A few times (less than half the time)
  □ Almost never or never
4. Over the past 4weeks, how would you rate your **level** of sexual arousal (“turn on”) during sexual activity or intercourse?
  □ No sexual activity
  □ Very high
  □ High
  □ Moderate
  □ Low
  □ Very low or none at all
5. Over the past 4weeks, how **confident** were you about becoming sexually aroused during sexual activity or intercourse?
  □ No sexual activity Very
  □ High confidence
  □ Moderate Confidence
  □ Low Confidence
  □ Very low or no confidence
  □
6. Over the past 4weeks, how **often** have you been satisfied with your arousal (excitement) during sexual activity or intercourse?
  □ No sexual activity
  □ Almost always or always
  □ Most times (more than half the time)
  □ Sometimes (about half the time)
  □ A few times (less than half the time)
  □ Almost never or never
7. Over the past 4weeks, how **often** did you become lubricated (“wet”) during sexual activity or intercourse?
  □ No sexual activity
  □ Almost always or always
  □ Most times (more than half the time)
  □ Sometimes (about half the time)
  □ A few times (less than half the time)
  □ Almost never or never
8. Over the past4weeks, how **difficult** was it to become lubricated (“wet”) during sexual activity or intercourse?
  □ No sexual activity
  □ Extremely difficult or impossible
  □ Very difficult
  □ Difficult
  □ Slightly difficult
  □ Not difficult
9. Over the past 4weeks, how often did you **maintain** your lubrication (“wetness”) until completion of sexual activity or intercourse?
  □ No sexual activity
  □ Almost always or always
  □ Most times (more than half the time)
  □ Sometimes (about half the time)
  □ A few times (less than half the time)
  □ Almost never or never
10. Over the past 4weeks, how **difficult** was it to maintain your lubrication (“wetness”) until completion of sexual activity or intercourse?
  □ No sexual activity
  □ Extremely difficult or impossible
  □ Very difficult
  □ Difficult
  □ Slightly difficult
  □ Not difficult
11. Over the past 4weeks,when you had sexual stimulation or intercourse, how **Often** did you reach orgasm (climax)?
  □ No sexual activity
  □ Almost always or always
  □ Most times (more than half the time)
  □ Sometimes (about half the time)
  □ A few times (less than half the time)
  □ Almost never or never
12. Over the past 4weeks,when you had sexual stimulation or intercourse, how **difficult** was it for you to reach orgasm (climax)?
  □ No sexual activity
  □ Extremely difficult or impossible
  □ Very difficult
  □ Difficult
  □ Slightly difficult
  □ Not difficult
13. Over the past 4weeks, how **satisfied** were you with your ability to reach orgasm (climax) during sexual activity or intercourse?
  □ No sexual activity
  □ Very satisfied
  □ Moderately satisfied
  □ About equally satisfied and dissatisfied
  □ Moderately dissatisfied
  □ Very dissatisfied
14. Over the past 4weeks,how **satisfied** have you been with the amount of emotional closeness during sexual activity between you and your partner?
  No sexual activity
  Very satisfied
  Moderately satisfied
  About equally satisfied and dissatisfied
  Moderately dissatisfied
  Very dissatisfied
15. Over the past 4weeks, how **satisfied** have you been with your sexual relationship with your partner?
  □ Very satisfied
  □ Moderately satisfied
  □ About equally satisfied and dissatisfied
  □ Moderately dissatisfied
  □ Very dissatisfied
16. Over the past 4weeks, how **satisfied** have you been with your overall sexual life?
  □ Very satisfied
  □ Moderately satisfied
  □ About equally satisfied and dissatisfied
  □ Moderately dissatisfied
  □ Very dissatisfied
17. Over the past 4weeks, how **often** did you experience discomfort or pain <u>during</u> vaginal penetration?
  □ Did not attempt intercourse
  □ Almost always or always
  □ Most times (more than half the time)
  □ Sometimes (about half the time)
  □ A few times (less than half the time)
  □ Almost never or never
18. Over the past 4weeks, how **often** did you experience discomfort or pain <u>following</u> vaginal penetration?
  □ Did not attempt intercourse
  □ Almost always or always
  □ Most times (more than half the time)
  □ Sometimes (about half the time)
  □ A few times (less than half the time)
  □ Almost never or never
19. Over the past 4weeks, how would you rate your **level** (degree) of discomfort or pain during or following vaginal penetration?
  □ Did not attempt intercourse
  □ Very high
  □ High
  □ Moderate
  □ Low
  □ Very low or none at all

***Thank you for completing this questionnaire***

Copyright □ 2000AllRightsReserved

## Appendix II

### CONSENTFORM

                            Serial number…………………..

Evaluation of female sexual function in type 2 diabetes mellitus patients in Umuahia with emphasis on its frequency and predictors

I,………………………………………………………………………………………of………… ………………………………………………………………………………………….hereby consent to participate in the study on Evaluation of female sexual function in type 2 diabetes mellitus patients in Umuahia with emphasis on its frequency and predictors

Dr…………………………………………………..has explained the nature of the study with its benefits and risks to me. I understand that the study is to be carried out solely for the purpose of Medical Research and I am willing to act as a volunteer for that purpose.

Date………………………………

Signature………………………….

Witness to signature………………..

I confirm that I have explained to you the purpose and nature of the study and the risks involved, including the fact that any refusal to participate will not in any way affect your normal care by me or any other member of this institution. All information obtained in this study is strictly confidential.

Date…………………………

Signature……………

